# Circadian clock neurons use activity-regulated gene expression for structural plasticity

**DOI:** 10.1101/2024.05.25.595887

**Authors:** Seana Lymer, Keyur Patel, Jennifer Lennon, Justin Blau

## Abstract

*Drosophila* s-LNv circadian pacemaker neurons show dramatic structural plasticity, with their projections expanded at dawn and then retracted by dusk. This predictable plasticity makes s-LNvs ideal to study molecular mechanisms of plasticity. Although s-LNv plasticity is controlled by their molecular clock, changing s-LNv excitability also regulates plasticity. Here, we tested the idea that s-LNvs use activity-regulated genes to control plasticity. We found that inducing expression of either of the activity-regulated transcription factors Hr38 or Sr (orthologs of mammalian Nr4a1 and Egr1) is sufficient to rapidly expand s-LNv projections. Conversely, transiently knocking down expression of either *Hr38* or *sr* blocks expansion of s-LNv projections at dawn. We show that Hr38 rapidly induces transcription of *sif,* which encodes a Rac1 GEF required for s-LNv plasticity rhythms. We conclude that the s-LNv molecular clock controls s-LNv excitability, which couples to an activity-regulated gene expression program to control s-LNv plasticity.

## Introduction

Neuronal plasticity is key for neuronal function, for example during memory formation (Whitlock *et al*, 2006). However, neuronal plasticity can be maladaptive in post-traumatic stress disorder and addiction (Bremner *et al*, 2008; Dani *et al*, 2001), and is likely misregulated in disorders such as ASD and schizophrenia (Ben-Shachar & Laifenfeld, 2004; Bourgeron, 2015). At the cellular level, plasticity can alter the connectivity between two neurons by changing excitability or by changing post-synaptic receptor density. More dramatically, structural plasticity can change the size of a synapse and can even connect previously unconnected neurons or break existing connections (Holtmaat & Svoboda, 2009). However, the detailed molecular mechanisms and layers of regulation that underlie plasticity are not fully understood, partly because it is often challenging to identify the neurons undergoing plasticity in intact brains under physiological conditions, especially in adult animals.

The small ventral lateral neuron (s-LNv) subset of circadian pacemaker neurons in *Drosophila* release the neuropeptide Pigment Dispersing Factor (PDF) into the dorsal brain where many of the other ∼140 clock neurons are located (Yasuyama & Meinertzhagen, 2010). PDF is required for circadian rhythms in constant darkness (DD) and for the morning peak of activity in light:dark (LD) cycles (Renn *et al*, 1999) . s-LNvs show plasticity at multiple levels: they change their excitability (Cao & Nitabach, 2008), the structure of their projections (Fernandez *et al*, 2008), and they likely make and break synaptic connections with downstream neurons (Gorostiza *et al*, 2014). s-LNvs cycle through all of these changes every 24hr even in DD, meaning that the predictable plasticity of s-LNvs does not require external signals and is intrinsic. Knowing which neurons are plastic and when they will change make s-LN_v_s excellent “model neurons” in which to study molecular mechanisms of plasticity. Furthermore, having only 4 s-LNvs per hemisphere makes visualization and quantification of their plasticity straightforward, and there are excellent tools to manipulate gene expression in adult s-LN_v_s with precise spatial and temporal control.

The termini of adult s-LNvs are expanded around dawn and retracted at dusk (Fernandez *et al*., 2008) (see Figure 1A). This rhythm is controlled by the circadian clock since it persists in DD and is lost in mutants that lack a functional circadian clock (Fernandez *et al*., 2008; Herrero *et al*, 2017). Altered expression of several genes has been shown to alter structural plasticity in s-LNvs, including the transcription factor Mef2 and the cell adhesion molecule Fas2 (Sivachenko *et al*, 2013), the matrix metalloproteases Mmp1 and Mmp2 (Gorostiza *et al*., 2014), and the Rho1 GTPase (Petsakou *et al*, 2015). Of these, only Rho1 has so far been shown to rapidly retract s-LNv projections in a similar timeframe to endogenous s-LNv plasticity. We previously showed that over-expressing Rho1 for 12hr from dusk to dawn prevented s-LNvs from expanding, and that Rho1 activity in s-LN_v_ projections is rhythmic, peaking at dusk as s-LNv projections retract (Petsakou *et al*., 2015). Rho1 retraction of s-LN_v_ projections and rhythmic Rho1 activity both require normal levels of the Guanine nucleotide Exchange Factor (GEF) *Puratrophin-like* (*Pura*), which we initially identified by its rhythmic clock-controlled expression in LNvs (Petsakou *et al*., 2015; Ruben *et al*, 2012). We also recently showed that the FMRP mRNA binding protein is required for s-LNv plasticity, and that FMRP overexpression for only 4 hours can regulate plasticity in s-LNvs (Gundermann *et al*, 2023). At least part of the effect of FMRP is by binding and likely reducing translation of *still life* (*sif*) (Gundermann *et al*., 2023), which encodes a Rac1 GEF (Sone *et al*, 1997). Thus s-LNv plasticity seems to be a balance between Rho1 and Rac1 activity, with Rho1 regulated by transcriptional regulation of *Pura*, and Rac1 regulated by post-translational regulation of *sif*.

**Figure 1:**
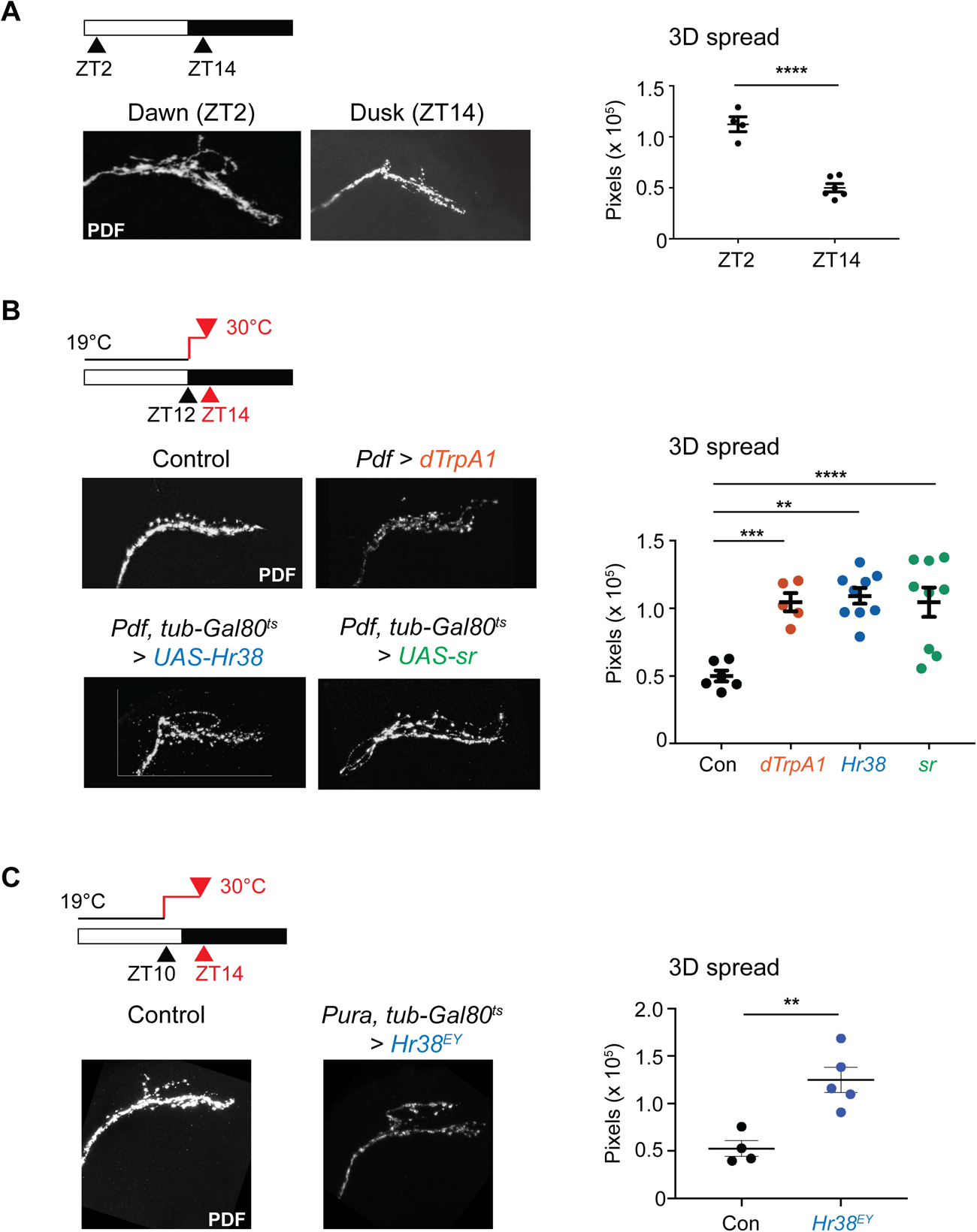
Expressing activity-regulated genes at dusk is sufficient to expand s-LNv projections. (A) **The normal cycle of s-LNv plasticity.** Control flies were kept in LD cycles at 25°C for at least 3 days, dissected at either ZT2 or ZT14 and stained with antibodies to PDF and then imaged. The 3D spread of the s-LNv projections was quantified using a MATLAB script. s-LNv projections have a larger 3D spread at ZT2 than at ZT14 (p < 0.001). Error bars show SEM. Statistics used student’s t-test. (B**) s-LN_v_ projections can be expanded at dusk by inducing s-LNvs to fire or by expressing *Hr38* or *sr* for 2 hours.** Flies were entrained to LD cycles at 19°C for at least 3 days and then shifted to 30°C to induce transgene expression for 2 hours starting at ZT12 (lights-off). Flies were dissected and fixed 2 hours later at ZT14. Left: Confocal images of s-LNv projections at ZT14 stained with anti-PDF. Top: Control (*Pdf-Gal4, tub-Gal80^ts^ > myrRFP*); TrpA1 (*Pdf-Gal4, tub-Gal80^ts^ > UAS-TrpA1*), Bottom: *Hr38* (*Pdf-Gal4, tub-Gal80^ts^ > UAS-Hr38*); *sr* (*Pdf-Gal4, tub-Gal80^ts^ > UAS-sr*) at ZT14. Right: Quantification of 3D spread of projections at ZT14 after inducing expression of control (*myrRFP*, black), *TrpA1* (orange), *Hr38* (blue) or *stripe* (*sr,* green) in LNvs. The data represent at least 5 brains from a minimum of 2 independent experiments. Error bars show SEM. Statistics used student’s t-test. ** = p < 0.01, *** = p < 0.001, **** = p < 0.0001. (C) **s-LNv projections can be expanded at dusk by inducing *Hr38* expression only in s-LNvs.** Flies were entrained to LD cycles at 19°C for at least 3 days and then shifted to 30°C. *Pura-Gal4*, *tub-Gal80^ts^* flies were crossed to either *y w* flies (control) or to *Hr38^EY^* flies, which have a P element containing UAS binding sites inserted in *Hr38* to induce *Hr38* expression. The temperature shift started at ZT10 and flies were dissected at ZT14. Left: Confocal images of s-LN_v_ projections at ZT14 stained with anti-PDF. Right: Quantification of 3D spread of s-LNv projections at ZT14. The data represent at least 4 brains per genotype. Error bars show SEM. Statistics used student’s t-test. ** = p < 0.01.

Making s-LNvs fire at dusk by activating an ectopically expressed cation channel is sufficient to expand their projections in only 2 hours (Sivachenko *et al*., 2013). The speed of this effect is reminiscent of activity-dependent gene expression (Yap & Greenberg, 2018). Furthermore, over-expressing the activity-dependent transcription factor Mef2 made s-LNvs constitutively expanded (Sivachenko *et al*., 2013), although this seems to require longer-term expression of Mef2 (Petsakou *et al*., 2015). The connections between s-LNv expansion and neuronal activity led us to test the role of activity-dependent gene expression in s-LNv structural plasticity.

Activity-dependent gene expression links neuronal activity to synaptic plasticity in mammalian neurons (Flavell & Greenberg, 2008; Yap & Greenberg, 2018). Transcription of a set of activity-regulated genes (ARGs), also known as immediate-early genes, increases within minutes of neuronal firing (Greenberg *et al*, 1986). Some ARGs such as Arc directly function in plasticity, while others encode transcription factors that regulate a second set of downstream genes, several of which are direct effectors of plasticity. ARGs and their downstream genes vary in different neurons, which makes identification of plasticity genes challenging because different neuronal subtypes are often analyzed together (Chen *et al*, 2016; Yap & Greenberg, 2018).

ARGs are a likely common response to neuronal firing across species e.g. (Fujita *et al*, 2013). Therefore, we decided to test if ARGs are important in s-LN_v_ expansion since s-LNv firing increases at dawn (Cao & Nitabach, 2008) at the same time of day as s-LNv projections expand (Fernandez *et al*., 2008). We found that over-expressing either of two ARGs – *Hormone receptor-like in 38* (*Hr38)* or *stripe* (*sr*) – is sufficient to drive s-LNv expansion at dusk when s-LN_v_s are normally retracted and are required for s-LNv structural plasticity at dawn. We found that transcription of *sif* increases at dawn and that *sif* transcription is rapidly upregulated by expressing *Hr38* at the wrong time of day. We conclude that *sif* is a key regulator of s-LNv plasticity and is regulated at two distinct levels: transcriptionally by Hr38, and post-transcriptionally by FMRP.

## Results

### Activity-regulated gene expression is sufficient to drive s-LNv axonal expansion

We first wanted to confirm that inducing neuronal activity at dusk is sufficient to expand s-LNv projections. For this, we used *Pdf-Gal4* and *tubulin-Gal80^ts^* (*tub-Gal80^ts^*) to temporally and spatially control expression of *UAS-TrpA1* in LNvs. The only other cells that produce PDF in the central brain are the large LNvs (l-LNvs), which are not important for circadian behavior (Stoleru *et al*, 2005). Thus *Pdf-Gal4* only expresses in the s- and l-LNvs, and we use it here extensively as a very specific central brain Gal4 driver (Park *et al*, 2000). *tub-Gal80^ts^* represses Gal4 activity at temperatures below 25°C, but is inactivated at higher temperatures (McGuire *et al*, 2003), which allows Gal4-activated transgene expression. *TrpA1* encodes a heat-activated *Drosophila* cation channel which can be ectopically expressed in neurons and induce firing in response to high temperature (Hamada *et al*, 2008).

We raised flies at 19°C to block Gal4 activity, and then shifted flies to 30°C to induce *TrpA1* expression and to simultaneously activate newly-synthesized TrpA1 to depolarize LNvs. Flies were entrained to 12 hour : 12 hour Light Dark (LD) cycles at 19°C for at least 3 days in this and all other temperature shift experiments, and then shifted to 30°C at Zeitgeber Time 12 (ZT12; lights on from ZT0-12; lights off from ZT12-24). Fly brains were dissected and fixed 2 hours later at ZT14. s-LNv projections were visualized by staining with antisera to the neuropeptide PDF, and we then used a MATLAB script to reconstruct s-LNv termini from confocal z-stacks and quantify their spread in three dimensions (Petsakou *et al*., 2015). We used flies with *Pdf-Gal4* and *tub-Gal80^ts^* controlling expression of *UAS-myrRFP* as control flies for this experiment and most of the other adult fly experiments. Control and experimental flies were entrained in the same LD cycles and experienced the same temperature shifts.

The data in Figure 1B show that inducing expression and activity of TrpA1 at dusk for 2 hours is sufficient to expand s-LNv projections to a dawn-like state (see also Figure 1A), consistent with previous data (Sivachenko *et al*., 2013). Control flies (*Pdf, tub-Gal80^ts^ > myrRFP*) have retracted s-LNv projections at dusk, verifying that shifting temperature alone does not expand s-LNv projections. Thus expansion of s-LNv projections is not a consequence of time of day *per se*, but rather of s-LNv neuronal activity. This leads to the idea that expansion of s-LNv projections follows a circadian rhythm *because* s-LNv neuronal activity is rhythmic and normally increases just prior to dawn (Cao & Nitabach, 2008) – as do intracellular Ca^2+^ levels (Liang *et al*, 2016).

Neuronal activity rapidly activates transcription of ARGs. We tested two ARGs that are expressed in LNvs: *Hormone receptor-like in 38* (*Hr38)* and *stripe* (*sr*) (Chen *et al*., 2016), both of which encode transcription factors. *Hr38* was the first ARG identified in flies (Fujita *et al*., 2013) and its orthologue *Nr4a1* is an ARG in mammals (Sheng & Greenberg, 1990). *Hr38* expression is rhythmic in adult s-LNvs, peaking at dawn when s-LNv firing increases (Kula-Eversole *et al*, 2010). We recently showed that an *Hr38* transcriptional reporter gene is rhythmic in larval LNvs (Zhu *et al*, 2024), which become the adult s-LNvs (Kaneko & Hall, 2000). Furthermore, we found that *Hr38* transcription increases when larval LNvs are induced to fire at dusk, showing that *Hr38* is a *bona fide* ARG in LNvs (Zhu *et al*., 2024). *sr* is a second ARG expressed in s-LNvs (Chen *et al*., 2016). *Egr1*, the mammalian Sr ortholog, is also activity-regulated and plays a role in neuronal plasticity and learning (James *et al*, 2005).

To test if expression of ARGs can drive expansion of s-LNv projections, we used *Pdf-Gal4, tub-Gal80^ts^* to control the timing of expression of transgenes for either *Hr38* (*UAS-Hr38*) or *sr* (*UAS-sr*). We induced expression of each transgene at ZT12 when projections are normally retracted, and measured s-LNv projections after 2 hours at 30°C as we had done with *UAS-TrpA1*. The data in Figure 1B show that a 2hr induction of either *Hr38* or *sr* was sufficient to expand s-LNv projections to a dawn-like state.

*Pdf-Gal4* is expressed in both s-LNvs and l-LNvs (Park *et al*., 2000). To confirm that *Hr38* acts cell-autonomously, we used the *Pura-Gal4* driver, which is expressed in s-LNvs but not l-LNvs (Sekiguchi *et al*, 2020). We also used an independent way to express *Hr38*: an EPgy2 P-element inserted ∼2kb upstream of the start site of *Hr38* transcription, which we refer to as *Hr38^EY^*. EPgy2 contains binding sites for Gal4 and thus this insertion should lead to *Hr38* overexpression in a Gal4-dependent manner, which we confirmed in Figure 3.

Flies were again raised until adulthood at 18°C and then entrained in LD cycles also at 18°C. We then shifted flies to 30°C for 4 hours starting at ZT10 before dissecting and fixing brains at ZT14. We compared the s-LNv projections in control and experimental flies. The control flies were the progeny of *Pura-Gal4, tub-Gal80^ts^* flies crossed to *y w* flies, and the experimental flies were the progeny of *Pura-Gal4, tub-Gal80^ts^*flies crossed to *Hr38^EY^* flies. The data in Figure 1C show that the experimental flies have expanded dawn-like s-LNv projections after 4 hours of inducing *Hr38* expression in s-LNvs, whereas the control flies had retracted s-LNv projections. Thus we conclude that activity-dependent transcription factors in s-LNvs are sufficient to induce expansion of their projections at dusk, when their expression is normally low (Chen *et al*., 2016; Kula-Eversole *et al*., 2010; Zhu *et al*., 2024).

### Activity-regulated gene expression is required for full expansion of s-LN_v_ projections

The data in Figure 1 shows that an activity-regulated gene expression program can drive expansion of s-LNv projections. To test if ARG expression is part of the normal program that s-LNvs use to expand their projections, we used transgenes to transiently block ARG expression at dawn. We again used *Pdf-Gal4* and *tub-Gal80^ts^* to control the timing of transgene expression - in this case *UAS-RNAi* transgenes that target either *Hr38* or *sr*. We found that inducing expression of either *Hr38-RNAi* or *sr-RNAi* at ZT12 blocks expansion of s-LNv projections 14 hours later at ZT2 with s-LNv projections in a retracted dusk-like state (Figure 2A).

**Figure 2:**
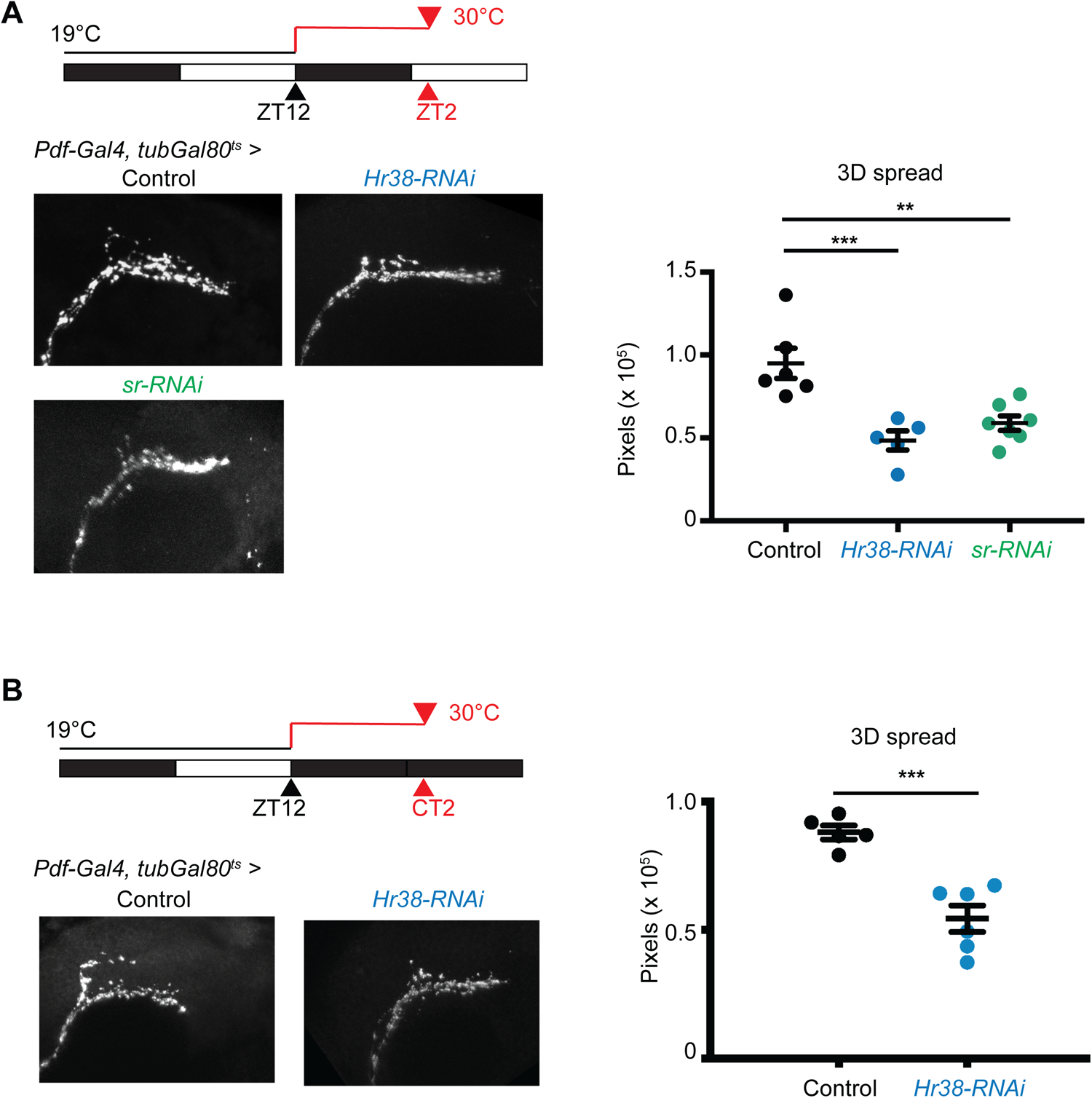
Activity-regulated genes are required to expand s-LNv projections at dawn. (A) Left: Confocal images of s-LNv projections maintained in LD cycles at 19°C and then shifted to 30°C starting at ZT12. Flies were dissected at ZT2 (2 hours after lights on) after 14 hours of transgene induction and stained with anti-PDF. Control (*Pdf-Gal4, tub-Gal80^ts^ > myrRFP*); *Hr38-RNAi* (*Pdf-Gal4, tub-Gal80^ts^ > Hr38-RNAi*); *sr-RNAi* (*Pdf-Gal4, tub-Gal80^ts^ > sr-RNAi*) Right: Quantification of 3D spread of projections at ZT2 after inducing *Hr38-RNAi* (blue) or *stripe-RNAi* (green) in LNvs. The data are an average of at least 5 brains from at least 2 independent experiments. The maximum projection image shown for *sr-RNAi* had brightness & contrast edits in ImageJ to improve visibility of the dorsal turn of the s-LNv projections. Min,Max (0,255) was changed to Min,Max(8,107.6). Error bars show SEM. Statistics used student’s t-test. ** = p < 0.01, *** = p < 0.001. (B) Left: Confocal images of s-LNv projections maintained in LD cycles at 19°C and then shifted to 30°C starting at ZT12. Flies were kept in DD and dissected at CT2 (2 hours after subjective lights on), and then stained with anti-PDF. Control (*Pdf-Gal4, tub-Gal80^ts^ > myrRFP*); *Hr38-RNAi* (*Pdf-Gal4, tub-Gal80^ts^ > Hr38-RNAi*). Right: Quantification of 3D spread of projections at CT2 after inducing *Hr38-RNAi* (blue) in LNvs. The data are an average of at least 5 brains from 1 experiment. Error bars show SEM. Statistics used student’s t-test. *** = p < 0.001.

We also tested if *Hr38* expression is required for s-LNv projections to expand in constant darkness (DD). Flies were entrained in LD at 19°C and transferred to 30°C at ZT12 to induce either a control transgene (*UAS-myrRFP*) or *UAS-Hr38-RNAi*. Flies were dissected 14 hours later at CT2 (Circadian Time, time in DD after entrainment to LD cycles), with flies maintained in darkness instead of returning to the light portion of an LD cycle. The data in Figure 2B show expanded s-LNv projections in control flies that went through this light and temperature regime which is expected because s-LNv plasticity persists in DD (Fernandez *et al*., 2008). However, inducing *Hr38-RNAi* in LNvs for 14 hours blocked the expansion of s-LNv projections. Thus *Hr38* is required for full expansion of s-LNv projections in both LD and DD. Furthermore, the data from DD indicate that the activity-dependent gene expression program likely occurs via circadian regulation of s-LNv neuronal activity and does not depend on light-induced firing (Yuan *et al*, 2011).

Together, the gain-of-function (Figure 1) and knockdown data (Figure 2) indicate that *Hr38* and *sr* are very important for s-LNv expansion at dawn. Figure 1 showed that expressing these activity-regulated genes at the wrong time of day is as effective as inducing firing at expanding s-LNv projections. Figure 2 shows that *Hr38* and *sr* expression are both required for s-LNv projections to expand at dawn. Thus we conclude that an activity-dependent gene expression program drives the dawn plasticity of s-LNv projections.

### Transcription of *still life* (*sif*) is upregulated by Hr38

Since both *Hr38* and *sr* encode transcription factors, we wanted to know what genes they regulate to promote expansion of s-LNv projections at dawn. In separate work, we have found that Sr represses transcription of the plasticity gene *Pura* (Zhu *et al*., 2024). We therefore focussed on Hr38 targets in this study.

One candidate Hr38 target is *still life* (*sif*). We have found that *sif* mRNA is bound by the RNA-binding protein FMRP in s-LNvs (Gundermann *et al*., 2023). FMRP is a bi-directional and rapid regulator of s-LNv plasticity: Flies with a null mutation in the *Fmr1* gene lose rhythms in s-LNv plasticity, while overproducing FMRP at dawn blocks the expansion of s-LNv projections (Gundermann *et al*., 2023). Finding *sif* as an FMRP target led us to test the importance of *sif* itself in expanding s-LNv projections. We also found that knocking down *sif* expression at dawn using RNAi is sufficient to block s-LNv projections from expanding (Gundermann *et al*., 2023), while overexpressing *sif* at dusk leads to expanded s-LNv projections. Thus appropriate levels of *sif* are clearly important in regulating s-LNv plasticity, and *sif* – like *Hr38* (Figure 2) – is required for s-LNv projections to expand at dawn (Gundermann *et al*., 2023).

We first wanted to know when *sif* is normally transcribed in LNvs. To measure this, we performed in situ hybridization with probes complementary to one of the large introns common to all *sif* isoforms. In situ hybridization with intronic probes labels nascent transcripts as 1-2 spots in the nucleus at the site of transcription (Yang *et al*, 2017). We also used sequences complementary to the large 4kb intron in *Hr38* to measure its transcription, and probes complementary to the single *Pdf* exon to label the cytoplasm of LN_v_s to be able to identify the relevant cells.

We decided to study the timing of *sif* transcription in larval rather than adult LNvs for four reasons. First, the rapid activation of activity-regulated genes in minutes makes it challenging to dissect sufficient adult brains in a short time. In contrast, larval brains are easy to rapidly dissect and thus the different brains in one sample would come from a time window of a few minutes.

Second, expression of the core clock genes is similar in larval and adult s-LNvs (Price *et al*, 1998). Indeed, the larval LNvs become the adult LNvs and even maintain their clock time from larvae to adults (Kaneko & Hall, 2000; Sehgal *et al*, 1992). Third, expression of an *Hr38* transcriptional reporter gene peaks shortly after dawn in larval LNvs (Zhu *et al*., 2024). Fourth, we identified how the plasticity gene *Pura* is regulated by studying larval LNvs, and then found that the *Pura* regulator Toy controls s-LNv plasticity in adult flies (Zhu *et al*., 2024). Thus although no plasticity rhythms have yet been described in larval LNvs, their gene expression patterns and timing are very similar to adult s-LNvs. In addition, we had previously found higher *sif* expression in larval LNvs isolated from *cycle* null mutants than *period* mutants, suggesting that *sif* expression is clock-regulated, although *sif* RNA rhythms have not been reported (Kula-Eversole *et al*., 2010; Ruben *et al*., 2012).

The data in Figure 3A show that neither *sif* nor *Hr38* transcription is detectable at ZT0, but both are clearly detectable one hour later at ZT1. *Hr38* transcription declines to low levels by ZT2. This pattern is expected for activity-regulated genes such as *c-fos* in mammalian neurons in culture whose transcription is rapidly activated before rapidly returning to low levels (Greenberg *et al*., 1986). The timing of *Hr38* transcription that we see here is consistent with data showing that *Hr38* mRNA levels rise in the *Drosophila* head in response to the end of a 30 minute alcohol exposure, peak at 1 hour and then return to steady state low levels by 2-3 hours (Adhikari *et al*, 2019).

**Figure 3:**
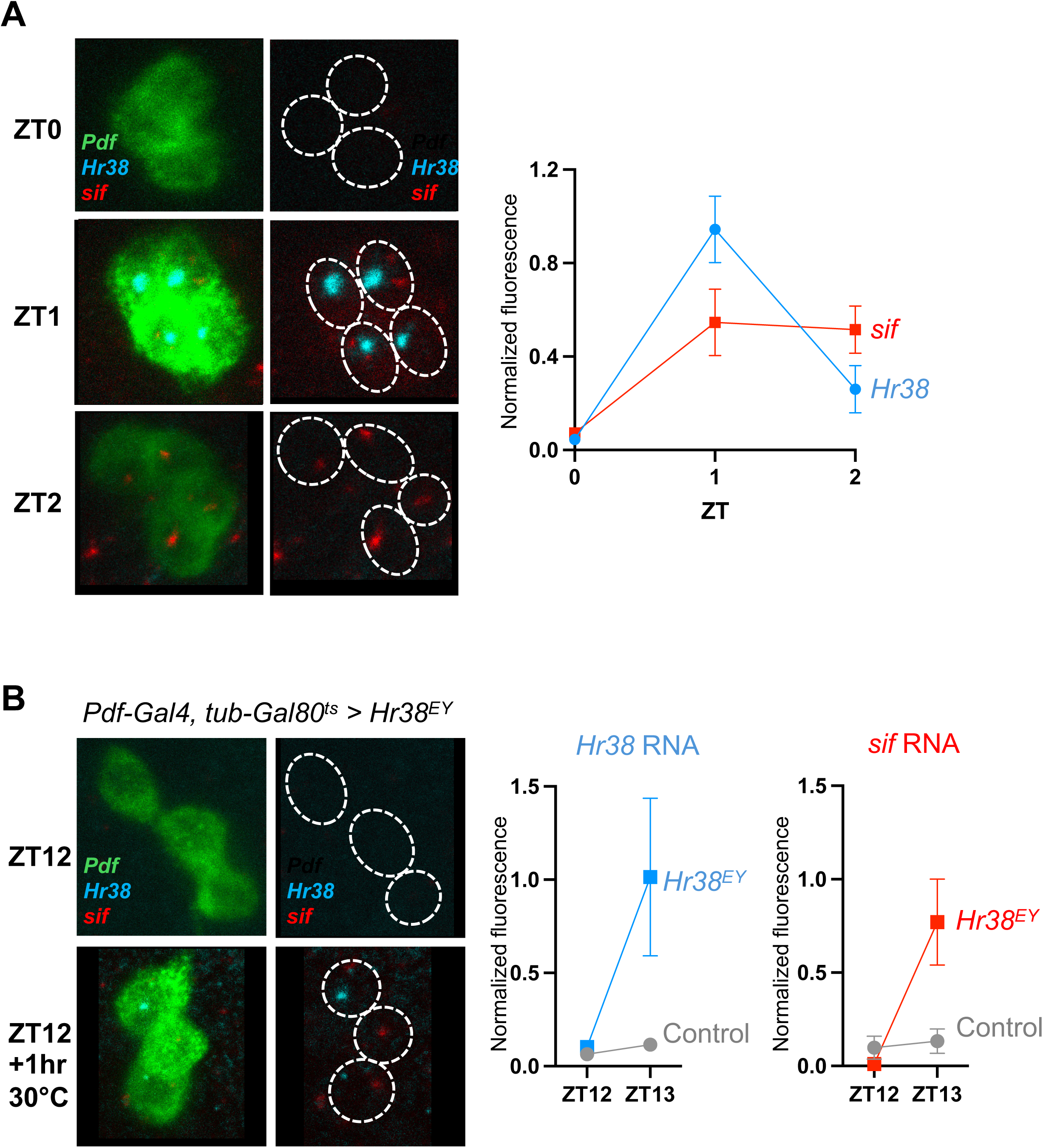
*sif* transcription increases at dawn and is rapidly upregulated by inducing *Hr38* expression. **(A) Time course of *Hr38* and *sif* transcription in larval LNvs.** Control *y w* larvae were dissected at ZT0, ZT1 or ZT2 for in situ hybridization with intronic probes for *Hr38* (blue) or *sif* (red) and exonic probes for *Pdf* (green). Images on the left show all 3 channels, while images on the right have the green *Pdf* signal replaced by a white dotted line to be able to see *Hr38* and *sif* intronic signals in the nucleus. Quantification on the right shows *Hr38* and *sif* levels relative to *Pdf*. **(B)** Inducing *Hr38* expression at dusk in LNvs rapidly increases *sif* transcription. *Pdf-Gal4, tubulin-Gal80^ts^* flies were crossed to either *y w* control flies, or *Hr38^EY^* flies. Larvae were raised at 18°C and entrained in LD cycles at 18°C. Larvae were dissected either at ZT12, or at ZT13 after raising the temperature to 30°C for 1 hour. In situ hybridization was then performed as above. The confocal images on the left show signals from intronic *Hr38* (blue) and *sif* probes (red), and exonic *Pdf* probes (green) applied to *Pdf-Gal4, tub-Gal80^ts^* > *Hr38^EY^* larvae. Images on the right show only the intronic probes, with white dotted lines replacing the *Pdf* signal. Quantification shows *Hr38* / *Pdf* or *sif* / *Pdf* fluorescence intensity for either control (grey line) or experimental (*Hr38^EY^*, colored line) larvae.

In contrast to *Hr38*, *sif* transcription was still detectable at ZT2 at similar levels to ZT1. These data are consistent with *sif* transcription being activity-dependent itself and/or dependent on *Hr38*, whose protein product would be expected to last for longer than the *Hr38* gene itself is transcribed.

To test if *sif* is regulated by *Hr38*, we took advantage of the *Hr38^EY^* insertion whose activation is sufficient to induce expansion of s-LNv projections at dawn (Figure 1C). We crossed flies with *Pdf-Gal4* and *tub-Gal80^ts^*to either *y w* control flies, or to *Hr38^EY^* flies as in Figure 1C. Larvae were raised at 18°C and entrained in LD cycles at 18°C. Larvae were then dissected at either ZT12, or at ZT13 having shifted the temperature in the incubator to 30°C beginning at ZT12. *Hr38* transcription is at low levels at these times in control LNvs (Figure 3B).

The data in Figure 3B show that neither *Hr38* nor *sif* transcription is detectable in larval LNvs at ZT12. However, LNvs with the *Hr38^EY^* insertion increased transcription of *Hr38* after one hour at 30°C at ZT13. This was specific to flies with the *Hr38^EY^* insertion as *Hr38* was not detected in control LNvs. These data confirm that the endogenous *Hr38* gene is induced in a Gal4-dependent manner.

We also detected *sif* expression at ZT13 in the experimental LNvs, but not in the control LNvs. Thus we conclude that Hr38 increases *sif* transcription. This experiment does not determine whether *sif* is a direct target of Hr38, although there cannot be many steps in between the Hr38 transcription factor and *sif* transcription given the speed of *sif* induction in this experiment and in wild type larvae in Figure 3A. We conclude that *sif* transcription is upregulated by Hr38, and thus that *sif* is part of the Hr38-mediated program that leads to expansion of s-LNv projections.

## Discussion

The *Drosophila* molecular clock drives daily rhythms in s-LNv neuronal excitability and plasticity (Cao & Nitabach, 2008; Fernandez *et al*., 2008). Our working model – see Figure 4 – is that the s-LNv molecular clock controls s-LNv excitability rhythms, and that increased firing of s-LNvs at dawn induces transcription of a set of activity-regulated genes, including *Hr38* and sr. This activity-regulated gene expression program then controls s-LNv structural plasticity with Hr38 up-regulating *sif* transcription (this paper) and Sr repressing *Pura* transcription (Zhu *et al*., 2024).

**Figure 4:**
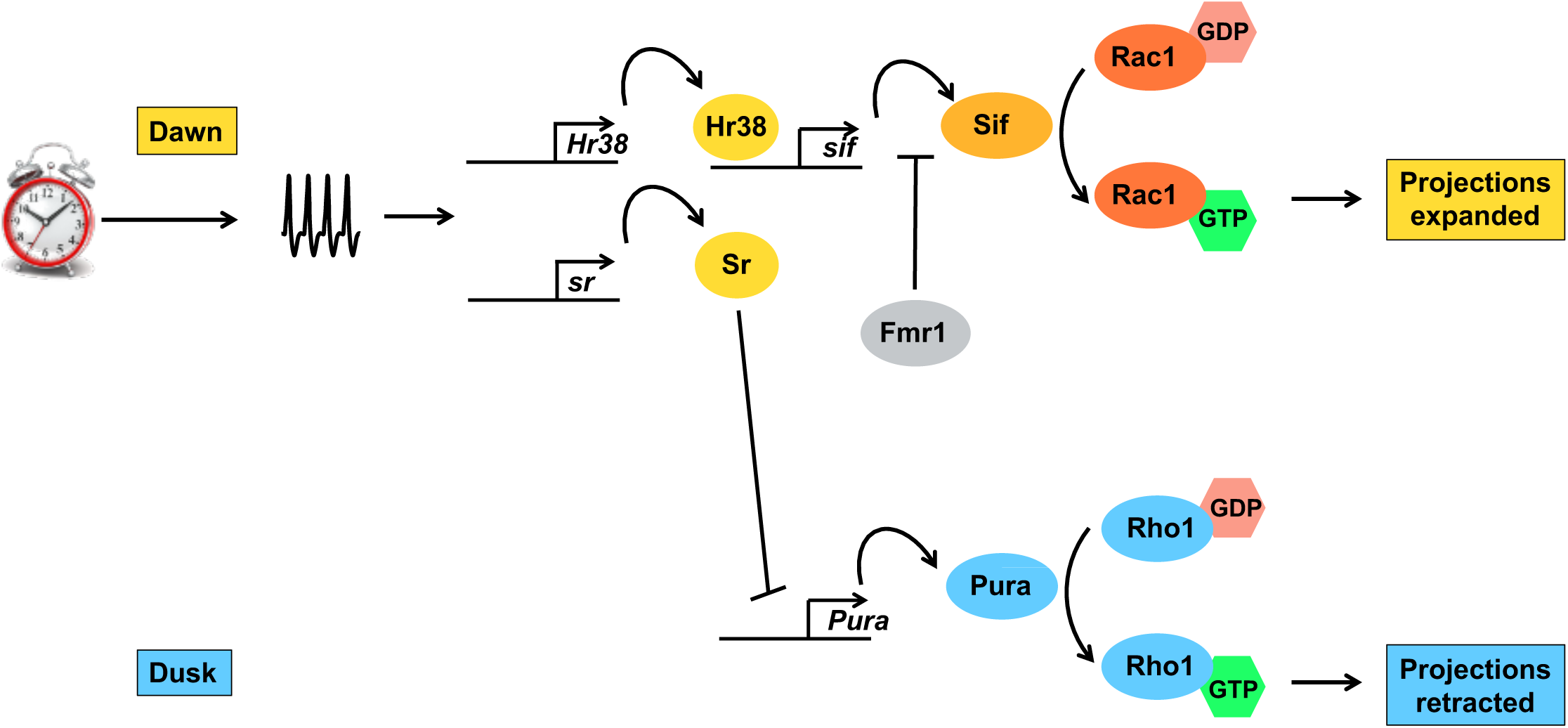
Model for how activity-dependent gene expression regulates s-LNv structural plasticity. The circadian molecular clock drives rhythmic excitability of s-LNv clock neurons that peak around dawn (Cao & Nitabach, 2008). *Hr38* and *sr* are activity-regulated genes in s-LN_v_s (Chen *et al*., 2016). Here we show that *Hr38* upregulates *sif* transcription, although we do not know if *sif* is a direct target of Hr38. *sif* and *Rac1* are required for s-LN_v_ plasticity at dawn, and *sif* is also a target of FMRP in s-LN_v_s, which likely blocks translation of *sif* mRNA (Gundermann *et al*., 2023). Sr represses transcription of *Pura* (Petsakou *et al*., 2015), a known plasticity gene.

This model explains how making s-LNvs fire at dusk rapidly changes the structure of their projections in a timeframe in which the gene expression profile of the circadian clock is unlikely to change dramatically. It also explains why clock mutants alter s-LNv plasticity (Fernandez *et al*., 2008; Herrero *et al*., 2017), because arguably the major function of the s-LNv clock beyond sustaining itself is to regulate s-LNv excitability. One way in which the s-LNv clock controls neuronal excitability is via rhythmic expression of *Irk1*, an inward-rectifier K+ channel (Ruben *et al*., 2012). In an analagous way, the DN1p clock neurons rhythmically express *Mid1*, also known as NCA localization factor 1 (Nlf-1), which helps localize the Narrow Abdomen Na+ channel to the membrane and control DN1p excitability (Flourakis *et al*, 2015). The same process may also occur in adult LNv clock neurons and in mammalian circadian pacemaker neurons in the suprachiasmatic nucleus (Flourakis *et al*., 2015).

Our data here on transcriptional regulation of *sif* adds to its layers of regulation in LNvs. We recently showed that *sif* is an FMRP target in s-LNvs (Gundermann *et al*., 2023). We also showed that the abnormally expanded s-LNv projections at dusk when *Fmr1* is knocked down can be rescued by simultaneously reducing *sif* expression. This led to the idea that FMRP normally represses *sif* mRNA translation at dusk in s-LNvs (Gundermann *et al*., 2023). *sif* is one of the very few genes that shows rhythms in alternative splicing in 3 different groups of clock neurons, including the LNvs (Wang *et al*, 2018), although this study did not distinguish between s-and l-LNvs. Thus *sif* is regulated at multiple levels in circadian neurons.

Sif is one of the few proteins that immunoprecipitated with the presynaptic protein Bruchpilot (Brp) in adult flies (Owald *et al*, 2010). Brp is located at active zones and is required to form T-bars (Fouquet *et al*, 2009; Kittel *et al*, 2006; Wagh *et al*, 2006) where neurotransmitter release takes place. Sif’s direct association with Brp is interesting because s-LNvs also undergo synaptic changes in addition to structural plasticity: s-LN_v_s have more pre-synaptic active zones at dawn and GRASP studies indicate that s-LN_v_s make and break connections with a 24 hour rhythm (Gorostiza *et al*., 2014). Indeed, Sif has a known role in synapse assembly at the *Drosophila* NMJ (Sone *et al*., 1997). Thus increased Sif levels at dawn may not just expand s-LNv projections, but also help build active synapses and connect s-LNvs to downstream target neurons.

The model in Figure 4 shows part of the transcriptional network logic that allows s-LNv projections to switch between their expanded state at dawn, and the retracted state at dusk. Increased s-LNv activity activates *sif* transcription and also represses *Pura* transcription. Ultimately it is probably the balance of Rac1 and Rho1 activity that determines the state of s-LNv projections and thus their ability to communicate with downstream neurons (Gundermann *et al*., 2023; Petsakou *et al*., 2015). While not all neurons may use this switch-type regulation, we note that the female hippocampus shows rhythms in dendritic spine number and synaptic density which follow the 4-5 day oestrous cycle (Jaric *et al*, 2019; Shors *et al*, 2001; Woolley & McEwen, 1992). The molecular mechanisms may even be similar with the mammalian Sr ortholog Egr1 linked to these changes in plasticity in female hippocampal neurons (Rocks *et al*, 2023).

The model in Figure 4 focuses on expression of *sif* and *Pura*, the GEFs that regulate Rac1 and Rho1 activity respectively in s-LNvs. There are likely to be other targets of Hr38 and Sr that play a role in structural plasticity. Indeed, a recent study showed that Nr4a1, one of the Hr38 orthologs in mammals, regulates expression of several cell adhesion molecules in inhibitory neurons in the mouse forebrain (Huang *et al*, 2024). It will therefore be interesting to find other Hr38 and Sr target genes and test how they function in s-LNv structural plasticity.

## Methods

### Fly strains

The following fly strains were used in this study: *Pdf-Gal4* (Park *et al*., 2000); *tubulin-Gal80^ts^* (McGuire *et al*., 2003); *UAS-Hr38* (FlyORF); *UAS-Hr38-RNAi* (Bloomington: 29376); *UAS-stripe* (Bloomington: 36533); *UAS-stripe-RNAi* (Bloomington: 60024); *UAS-TrpA1* (Bloomington: 26264) (Hamada *et al*., 2008); *Hr38^EY14161^*, referred to as *Hr38^EY^* in the text (Bloomington: 20910). All crosses were raised at 18°C because of the *tubulin-Gal80^ts^*transgene.

### Immunofluorescence

Immunofluorescence was performed as in (Pinto-Teixeira *et al*, 2018). Brains were dissected in PBS (phosphate-buffered saline) and fixed in 4% formaldehyde for 20 minutes at room temperature. Brains were rinsed three times in PBT (PBS with 0.3% Triton) including a final wash of at least an hour. Brains were incubated in primary antibody at least overnight at 4°C, then washed three times in PBT and incubated with secondary antibody either overnight at 4°C or for 1-2 hour at room temperature. Brains were rinsed three times and left to wash overnight before mounting in Slowfade.

#### Antibodies used

PDF: “PDF C7”, Developmental Studies Hybridoma Bank (mouse, used at 1:50); Alexa Fluor 488 or 647, Life Technologies (donkey anti-mouse, used at 1:500).

### Imaging and quantification of structural plasticity

To quantify structural plasticity, projections were imaged and processed as in (Petsakou *et al*., 2015). Brains were scanned on a Leica SP5 or SP8 confocal microscope. Confocal z-stack images were imported to MATLAB and axonal projections were manually selected starting from where the s-LNv axons turn dorsally. The MATLAB script calculates the average spread in x, y, and z axes, and the product of these values is the 3D spread. All confocal images shown in the manuscript are maximum projections generated via ImageJ, converted to 32-bit grayscale.

### In situ hybridization

We used HCR probes designed and synthesized by Molecular Instruments and a protocol similar to (Herre *et al*, 2022). Probes recognized the single *Pdf* exon, the first intron of *Hr38* or the second intron of *sif.* Larval brains were dissected in PBS and fixed in 4% PFA for 20 minutes and then washed in PBST (0.1% Tween). Brains were dehydrated with a MeOH series of increasing concentrations on ice and stored in 100% MeOH at -20°C. The following steps were then performed on ice: Brains were rehydrated in a MeOH series of decreasing concentrations and washed twice in PBST before permeabilizing in 5% acetic acid for 5 minutes and washing in PBST. Brains were refixed in 4% PFA for 20 minutes, and then washed in PBST, followed by a 1:1 mix of PBST and 5x SSCT, and then 2 washes in 5x SSCT. Brains were then incubated in Hybridization solution (Molecular Instruments) for 5 mins at room temperature and then in fresh hybridization solution for 30 mins at 37°C. 0.4 μL of each probe was added to brains in 100 μL hybridization solution over two nights at 37°C.

Hybridization solution was then removed from samples which were washed 5 times in Wash buffer (Molecular Instruments) at 37°C, and then twice at room temperature in 5X SSCT. Brains were then equilibrated in Amplification buffer (Molecular Instruments) for 5 minutes.

Amplification probes were prepared according to the manufacturer’s instructions with 1.5 μl of the relevant amplification probes added to brains in 50 μl of Amplification buffer. Samples were then left for 5 hours. Samples were then washed in 5x SSCT at room temperature, brains mounted in SlowFade Gold and then imaged on a Leica SP8 confocal microscope.

## Acknowledgements

We are indebted to numerous stock centers for sharing flies and antibodies: the Bloomington *Drosophila* Stock Center (supported by NIH P40OD018537), the Developmental Studies Hybridoma Bank (created by the NICHD and maintained at The University of Iowa), the TRiP stock center at Harvard Medical School (supported by NIGMS, NIH) and FlyORF (Bischof *et al*, 2013). We thank Simon Kidd for bringing HCR in situ hybridization to the lab and for his earlier work on this project and to David Owald for his insightful comments on Sif function at the NMJ. This investigation was conducted in facilities constructed with support from the NIH National Center for Research Resources. SL was partly supported by the NYU’s Graduate School of Arts and Science MacCracken Program. KP was supported by an NYU CAS Dean’s Undergraduate Research Fellowship. This work was supported by NIH grant GM136363 (JB).

